# Lepidopteran Synteny Units (LSUs) reveal deep conservation of macrosynteny in butterflies and moths

**DOI:** 10.1101/2023.03.22.533864

**Authors:** Walther Traut, Ken Sahara, Richard H. ffrench-Constant

**Affiliations:** Institut für Biologie, Zentrum für Medizinische Struktur- und Zellbiologie, Universität zu Lübeck, Ratzeburger Allee 160, D-23562 Lübeck, Germany; Laboratory of Molecular Entomology, Faculty of Agriculture, Iwate University, 3-18-8, Ueda, Morioka, 020-8550, Japan; Centre for Ecology and Conservation, University of Exeter, Penryn Campus, Penryn, United Kingdom

**Keywords:** Chromosome evolution, macrosynteny, Lepidoptera, chromosome numbers, chromosome rearrangement, BUSCO

## Abstract

Advances in DNA sequencing technologies have, for the first time, provided us with enough whole chromosome-level genomes to understand in detail how chromosome number and composition change over time. Here, we use the genomes of butterflies and moths to look at the levels and age of macrosynteny in the Lepidoptera and Trichoptera. We used comparative BUSCO analsysis to define reproducible units of macrosynteny which we term ‘Lepidopteran Synteny Units’ or LSUs. The 31 chromosomes of the model butterfly *Melitaea cinxia* served as a reference point. The results show that chromosome-wide macrosynteny extends from the most basal branches of the Lepidopteran phylogeny to the most distal. This synteny also extends to the order Trichoptera, a sister group of the Lepidoptera. Thus, chromosome-wide macrosynteny has been conserved for a period of >200 My in this group of insects. We found no major interchromosomal translocations, reciprocal or non-reciprocal, in the genomes studied. Intrachromosomal rearrangements, in contrast, were abundant. Beyond its use in defining LSUs, this type of homology-based analysis will be useful in determining the relationships between chromosomal elements in different animals and plants. Further, by more precisely defining the breakpoints of chromosomal rearrangements we can begin to look at their potential roles in chromosomal evolution.

**Statement:** The authors declare no conflicting interests

**Contributions:** Conceptualisation: W.T., R.H.f.; data analysis: W.T.; writing & editing: W.T., K.S., R.H.f All authors read and approved the final manuscript.

## Introduction

Chromosome number varies considerably in both animals and plants, from n = 1 in the nematode *Parascaris univalens* (Boveri 1899; Goday and Pimpinelli 1986) and the ant *Myrmecia pilosula* (Crosland and Crozier 1986) to n = 720 in the fern *Ophioglossum reticulatum* (Khandelwal 2008). In animals, chromosome number and variability also differ between taxonomic groups. Within the insects, for example, the Diptera (the flies) have small numbers of chromosomes that show little variation, whereas the Orthoptera (the grasshoppers) have higher numbers of chromosomes. It is not clear what rules might govern chromosomal evolution between groups of species. There is for example no clear overall correlation between total genomic DNA content and chromosome number. Evolution of chromosome numbers might be driven by the need to link favourable combinations genes in synteny. - Here we use the term ‘synteny’ in its originally derived sense, the physical equivalent of the genetic term ‘linkage’ and disregarding gene order. - This hypothesis is referring to chromosomal territories in the interphase nucleus (Cremer and Cremer 2001, Meaburn and Misteli 2007) such genes are in close contact with each other.

To examine any potential biological role of chromosome diversity in genome evolution, we need to trace the ancestry of individual chromosomes. Here we develop a homology-based technique for exploring conserved synteny in the butterflies and moths. The Lepidoptera have previously been shown to have wide-ranging synteny or macrosynteny across autosomes and Z chromosomes via a range of different mapping techniques including comparative linkage mapping (Pringle et al. 2007; Beldade et al. 2009; Baxter et al. 2011; Van’t Hof et al. 2013), BAC-FISH mapping (Yasukochi et al. 2009; Yoshido et al. 2011; Sahara et al. 2013) by a comparison of annotated long-read sequences (d’Alençon et al.2010; The Heliconius Genome Consortium 2012), and by whole-genome comparison (Höök et al. 2023, Pazhenkova and Lukhtanov 2023).

Currently, the genomes of more than 200 different lepidopteran species have been fully sequenced, mainly by the Darwin Tree of Life Consortium (https://www.darwintreeoflife.org). These complete insect genomes therefore now afford a deeper analysis of macrosyteny across the different phylogenetic branches of the butterflies and moths. We selected 13 of these complete genomes representing species from different branches of the lepidopteran phylogeny plus one species of the sister order Trichoptera. To compare synteny across distantly related species we cannot rely on direct comparison of chromosomes and genomes, for example by dotplot analysis (Krumsiek et al. 2007). Instead, we used Benchmarking Universal Single-Copy Orthologs or BUSCO (Manni et al. 2021).

Such BUSCO analysis is classically used in genomics to determine the apparent completeness of a genome by comparing the observed versus the expected number of protein coding sequences predicted. However, here we use BUSCO derived scores containing the positional information on each BUSCO marker to compare the location of these markers in the chromosomes of different species. In this respect, the 5,286 BUSCO gene markers in the lineage-specific collection ‘lepidoptera_odb10’ present an unprecedented density of markers for all lepidopteran genomes. This comparative BUSCO analysis allows us to compare genomes from the most basal phylogenetic branch of the Lepidoptera to the more derived ones and even to a representative of a different insect order, the Trichoptera. Here, we show that butterfly and moth chromosomes are made up of well conserved synteny units, which we term Lepidopteran Synteny Units or LSUs. These LSUs are conserved in a wide phylogenetic spectrum of species including the basal group of Micropterigidae and the representative of the sister group Trichoptera. This means that the genome of ancestral Lepidoptera in the late Triassic period of geology was already organised into these basic macrosynteny units. We argue that such a homology-based system advances our understanding of how chromosomes evolve and that such a system avoids the confusion generated by simply comparing chromosome numbers.

## Materials and Methods Genomes and markers

We examined 13 lepidopteran genomes and one trichopteran genome (Table S1) chosen from across the Lepidopteran phylogenetic tree. We used the model species *Meltiaea cinxia* as a reference point for inter-species comparisons as it has a well characterized genome with 31 chromosomes. We located the chromosomal position of the 5,286 BUSCO markers from the dataset lepidoptera_odb10 (https://busco.ezlab.org/list_of_lineages.html) with BUSCO version 5.4.3 (https://busco.ezlab.org/) using default parameters. The “full_table.tsv” ouput of the program package lists the positional information of all markers from that set. We sorted the output from a BUSCO analysis of *Melitaea cinxia* chromosome-wise, and in a second round chromosome position-wise. Lines noted “Missing” or “Duplicated” were discarded. The chromosome-specific lepidoptera_odb10 marker subsets define the 31 LSUs. LSU_31 is the Z chromosome-specific subset of markers. Table S2 presents the list of all LSU markers ordered according to LSU and chromosome position.

### Phylogenetic analysis

In order to compare LSU markers across the Lepidopteran phylogeny, the BUSCO output from each species was combined with the LSU marker sets. Each BUSCO output line was assigned to the corresponding LSU marker line with ‘BUSCO_to_LSUs’ (Java code available on request). A line was deleted when the BUSCO output was “Missing” or “Duplicated”.

Therefore, all species comparisons are strictly based on single-copy orthologues. As the LSU markers were already arranged according to chromosome and chromosome position the succession of positional information from the BUSCO output lines reveals conservation and rearrangements in the compared chromosomes. Detection and counting of breakpoints was performed with ‘LSU_breakpoints’ (Java code available on request).

## Results

### Marker coverage

We chose the model butterfly *Melitaea cinxia* as an anchor genome as it has both a published chromosome-scale genome (Vila et al. 2021) and the assumed ancestral chromosome number of Lepidoptera, where n = 31. *M cinxia* chromosomes have previously been shown to have conserved synteny with chromosomes of other lepidopteran species (Ahola et al. 2014; Mongue et al. 2017; Traut et al. 2017). However the extent of this synteny across the Lepidoptera as a whole has not been examined. Here, using BUSCO (Manni et al.2021) we mapped 5210 of the 5286 markers from the lepidoptera_odb10 set to the *M. cinxia* genome. These markers covered most of the length of all chromosomes (Table 1). The marker subsets specific for the 31 chromosomes, autosomes plus Z chromosome, were then used to investigate the extent of conserved synteny in the other selected species. They exposed a similar breadth of coverage and well-conserved blocks of synteny which we here term Lepidopteran Synteny Units or LSUs. Table S2 lists the respective 31 LSU marker subsets of lepidoptera_odb10.

**Table 1.**
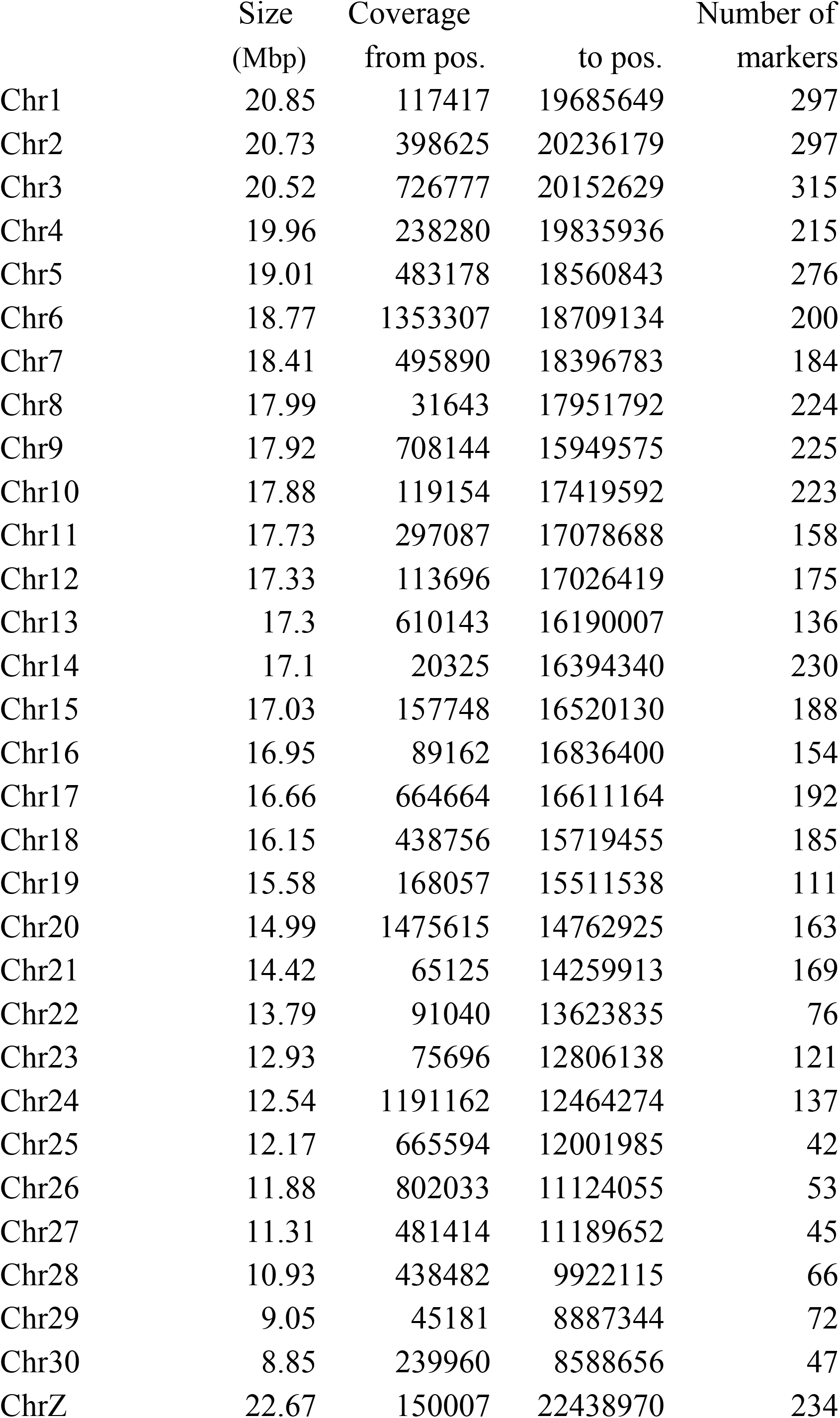
Coverage of *Melitaea cinxia* chromosomes with BUSCO markers

|  | Size<br>(Mbp) | Coverage<br>from pos. | to pos. | Number of<br>markers |
| --- | --- | --- | --- | --- |
| Chr1 | 20.85 | 117417 | 19685649 | 297 |
| Chr2 | 20.73 | 398625 | 20236179 | 297 |
| Chr3 | 20.52 | 726777 | 20152629 | 315 |
| Chr4 | 19.96 | 238280 | 19835936 | 215 |
| Chr5 | 19.01 | 483178 | 18560843 | 276 |
| Chr6 | 18.77 | 1353307 | 18709134 | 200 |
| Chr7 | 18.41 | 495890 | 18396783 | 184 |
| Chr8 | 17.99 | 31643 | 17951792 | 224 |
| Chr9 | 17.92 | 708144 | 15949575 | 225 |
| Chr10 | 17.88 | 119154 | 17419592 | 223 |
| Chr11 | 17.73 | 297087 | 17078688 | 158 |
| Chr12 | 17.33 | 113696 | 17026419 | 175 |
| Chr13 | 17.3 | 610143 | 16190007 | 136 |
| Chr14 | 17.1 | 20325 | 16394340 | 230 |
| Chr15 | 17.03 | 157748 | 16520130 | 188 |
| Chr16 | 16.95 | 89162 | 16836400 | 154 |
| Chr17 | 16.66 | 664664 | 16611164 | 192 |
| Chr18 | 16.15 | 438756 | 15719455 | 185 |
| Chr19 | 15.58 | 168057 | 15511538 | 111 |
| Chr20 | 14.99 | 1475615 | 14762925 | 163 |
| Chr21 | 14.42 | 65125 | 14259913 | 169 |
| Chr22 | 13.79 | 91040 | 13623835 | 76 |
| Chr23 | 12.93 | 75696 | 12806138 | 121 |
| Chr24 | 12.54 | 1191162 | 12464274 | 137 |
| Chr25 | 12.17 | 665594 | 12001985 | 42 |
| Chr26 | 11.88 | 802033 | 11124055 | 53 |
| Chr27 | 11.31 | 481414 | 11189652 | 45 |
| Chr28 | 10.93 | 438482 | 9922115 | 66 |
| Chr29 | 9.05 | 45181 | 8887344 | 72 |
| Chr30 | 8.85 | 239960 | 8588656 | 47 |
| ChrZ | 22.67 | 150007 | 22438970 | 234 |

### LSUs and chromosome numbers

For phylogenetic analysis we selected 12 other Lepidoptera and one Trichopteran. The species were chosen among those with available chromosome-scale genomes to represent different branches of the lepidopteran phylogeny (Figure 1). BUSCO searches in the selected genomes revealed nearly chromosome-sized blocks of conserved syntenic markers Lepidoptera-wide and even extending to Trichoptera (Table S3). Chromosome-wide macrosynteny (irrespective of marker gene order) is almost perfect among the chromosomes of the 12 heteroneuran Lepidoptera species examined here. These 12 species are from 11 different families: Nymphalidae, Lycaenidae, Noctuidae, Geometridae, Bombycidae, Blastobasidae, Zygaenidae, Cossidae, Plutellidae, Yponomeutidae, and Adelidae and therefore represent a broad cross-section of the clade Heteroneura. This clade comprises more than 99% of the ∼160,000 lepidopteran species (Kristensen 1999).

**Figure 1.**
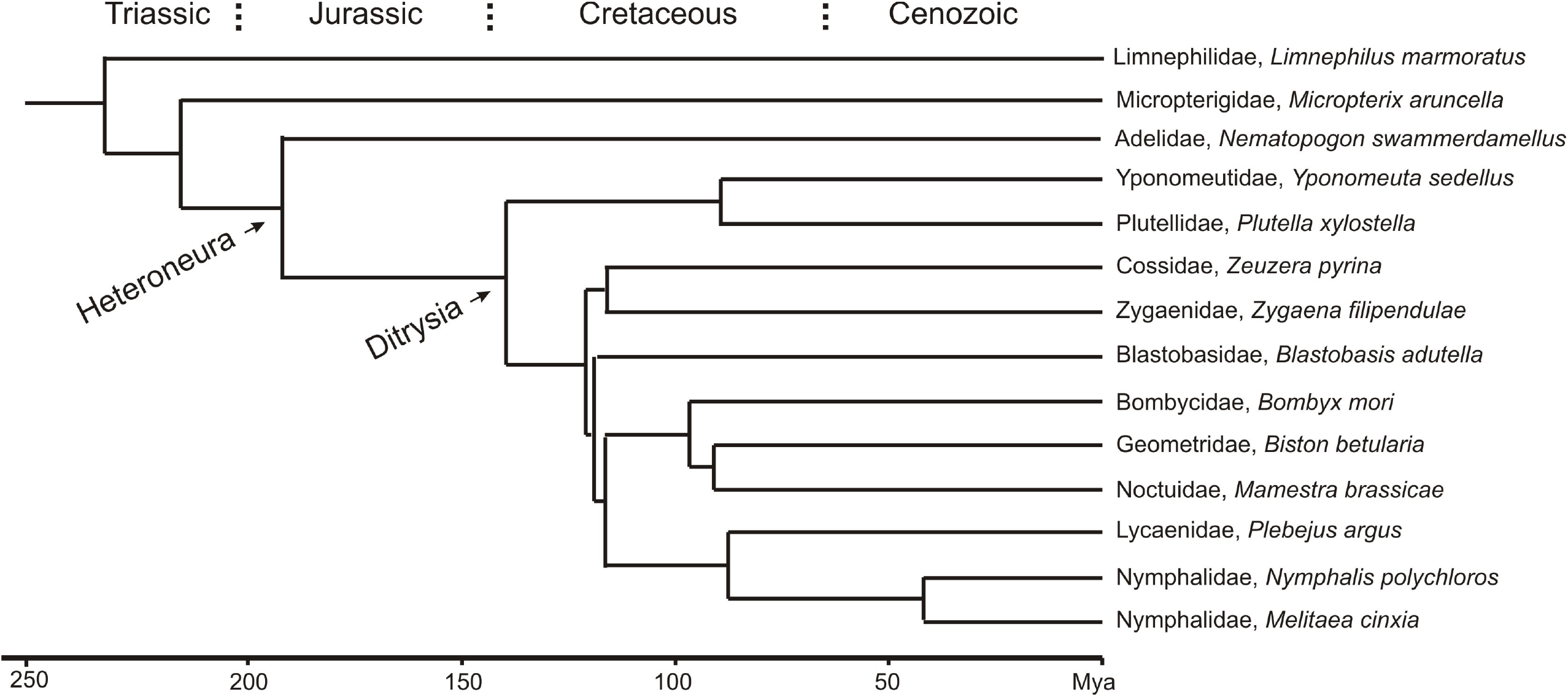
Phylogeny of the 13 lepidopteran species and the trichopteran species *Limnephilus marmoratus* studied in this paper. The phylogeny is based on data from Malm et al. (2013), Wahlberg et al. (2013), and Chazot et al. (2019).

Haploid chromosome numbers (autosomes plus Z chromosome) range from n = 23 to n = 31 among the heteroneuran species examined here. The representatives of the families Nymphalidae (*Melitaea cinxia* and *Nymphalis polychloros*), Noctuidae (*Mamestra brassicae*), Geometridae (*Biston betularia*), Plutellidae (*Plutella xylostella*), Cossidae (*Zeuzera pyrina*), Yponomeutidae (*Yponomeuta sedellus*), and Adelidae (*Nematopogon swammerdamellus*) have sets of n = 31 chromosomes. Each of the chromosomes constitutes one of the 31 LSUs defined here (Table S3). The representatives of the four remaining heteroneuran families have less than 31 chromosomes in their haploid sets of chromosomes. *Zygaena filipendulae* from the family Zygaenidae has n = 30 chromosomes. Table S3 shows that its chromosome 21 is a fusion product of two LSUs, LSU_25 and LSU_27. In *Blastobasis adustella* from the family Blastobasidae, another species with n = 30 chromosomes, it is the Z chromosome that is a fusion product of two LSUs, in this case LSU_21 and the original Z chromosomal unit LSU_31. The silkworm *Bombyx mori* from the Bombycidae family has n = 28 chromosomes. Three of the chromosomes, chromosomes 11, 23, and 24 can be seen composed of two LSUs each (Figure 2), confirming previous results (Yasukochi et al. 2016). The n = 23 chromosomes of *Plebejus argus* from the family Lycaenidae are the result of even more fusions, chromosome 1 consists of three LSUs, chromosomes 2 - 7 of two LSUs each (Figure 3). In summary, the 31 LSUs or 31 blocks of conserved synteny are an ancestral character in the clade of Heteroneura represented by those families. The situation is slightly different in *M. aruncella* and *L. marmoratus*, as discussed below.

**Figure 2.**
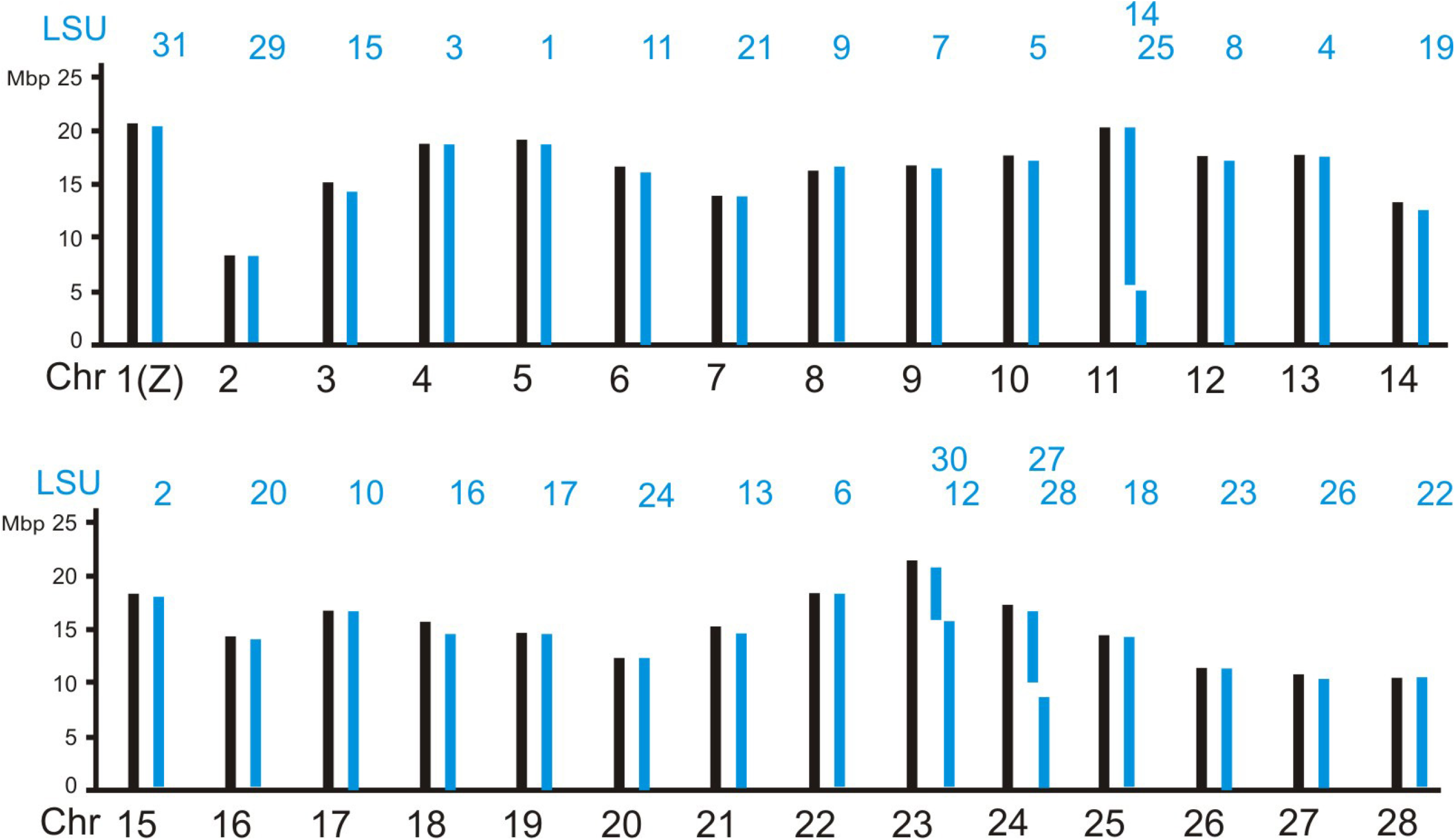
LSU composition of the 28 chromosomes from *Bombyx mori* (Bombycidae).

**Figure 3.**
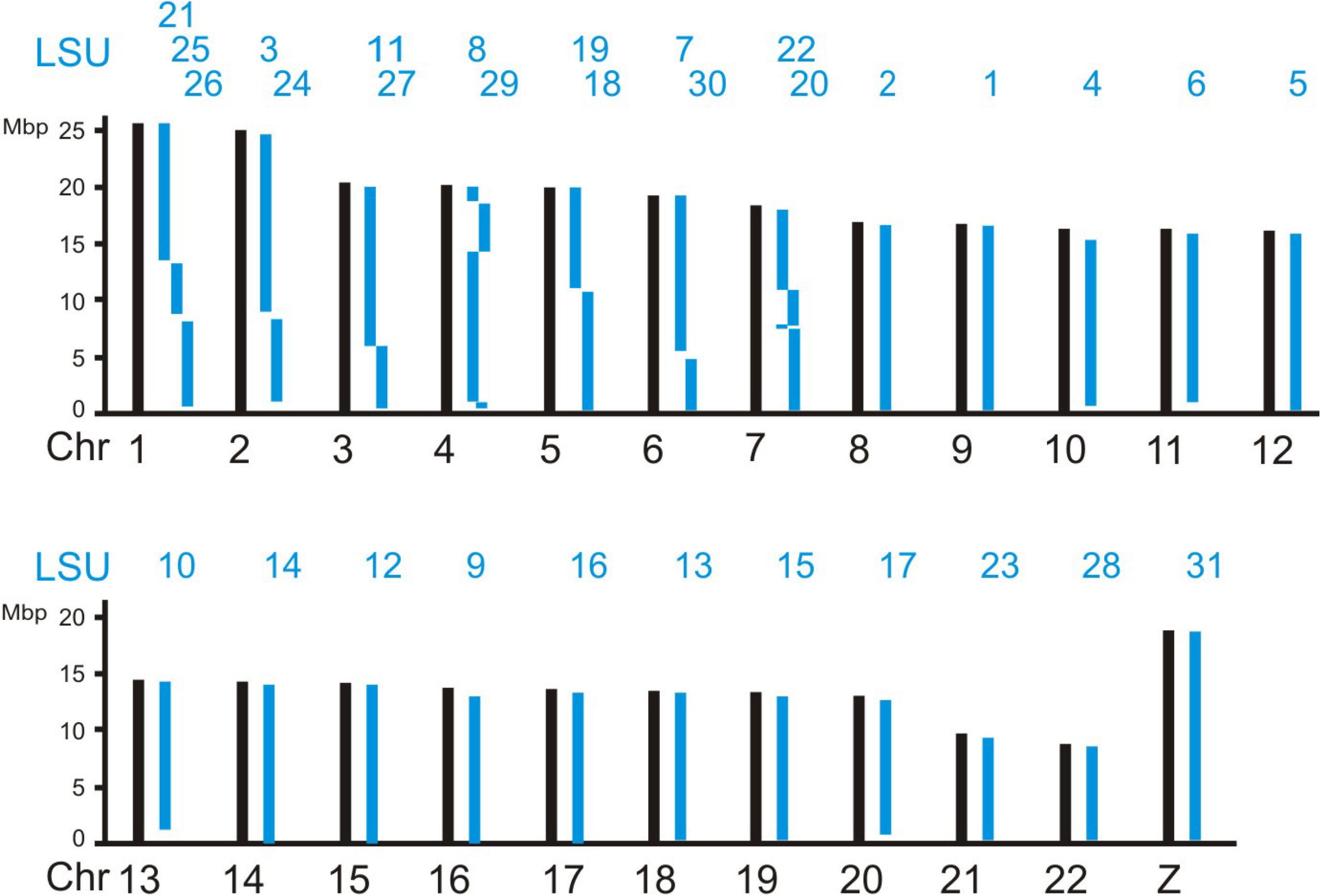
LSU composition of the 23 chromosomes from *Plebejus argus* (Lycaenidae).

### Chromosomal rearrangements

Having detected deep synteny across the Lepidoptera we then wanted to examine the frequency of inter- and intra-chromosomal rearrangements, such as inversions and translocations, via changes in the observed order of LSU markers. We selected four species with different divergence times for a study of such rearrangements, namely *N. polychlorus, B. mori, Y. sedellus*, and *N. swammerdamellus*, each compared with *M. cinxia*. No large inter-chromosomal translocations, either reciprocal or non-reciprocal, were detected. Small non-reciprocal transpositions of single or 2-3 adjacent markers happened at a low frequency of 0.01 - 0.27 per Mbp, 4 - 135 events per genome and, as expected, increased with divergence time (Table 2). In contrast intra-chromosomal rearrangements, however, were frequent. For a measure, we counted the breakpoints in the order of markers. Inversions leave two breakpoints, and translocations three breakpoints, as footprints in the rearranged chromosome. As expected, the number of breakpoints also increased with divergence time, from 96 to 1665 per genome. This corresponds to a rate of 0.46 - 3.34 per Mbp. It is important to note, however, that the resolution of rearrangement detection is limited by the density of available markers. The actual rate of rearrangements is presumably therefore much higher. For example, this analysis looses all rearrangements that can only be seen at the nucleotide level, such as those detected by comparisons of Pacbio sequences from closely related species (d’Alençon et al. 2010).

**Table 2.** Intra- and interchromosomal rearrangements detected as rearrangements within and between LSUs. Rearrangements registered in 4 species relative to *M. cinxia*. Time of divergence according to Wahlberg et al.(2013) and Chazot et al.(2019).

| Species | Breakpoints |  | Transposition of LSU markers |  |  | Time of divergence |
| --- | --- | --- | --- | --- | --- | --- |
|  | within LSUs | per Mbp | single | groups of 2 - 3 | per Mbp |  |
| *N. polychloros* | 232 | 0.46 | 4 | 0 | 0.01 | ~42 My |
| *B. mori* | 468 | 0.94 | 15 | 1 | 0.03 | ~116 My |
| *Y. sedellus* | 639 | 1.28 | 32 | 2 | 0.07 | ~139 My |
| *N. swammerdamellus* | 1665 | 3.34 | 135 | 1 | 0.27 | ~172 My |

#### *Micropterix aruncella* and *Limnephilus marmoratus as basal entities*

Lepidopteran macrosynteny extends to, but is less stringently conserved in, the more distantly related species *M. aruncella* from the basal lepidopteran family Micropterigidae and in *L. marmoratus* from the family Limnephilidae of the sister order Trichoptera (Table S3). Fully conserved are 30 of the 31 LSUs in at least one of the two species, and 24 still intact in the trichopteran *L. marmoratus*. But more genes are transferred to other chromosomes and a few larger interchromosomal rearrangements were documented by the mapped LSU markers. One of the synteny blocks, LSU_2, is well conserved in all heteroneuran species but is split in *M. aruncella* and *L. marmoratus*. This means in fact, that LSU_2 is a fusion product of two chromosomes. The fusion must have taken place during evolution of Lepidoptera after the branching off of Micropterigidae and before diversification of the Heteroneura. The other interchromosomal rearrangements may have taken place in the respective lineages. LSU_10 is entire in *M. aruncella* but split in *L. marmoratus* where it forms Chr27 and part of Chr10. In *L. marmoratus*, LSU_29 and LSU_30 together form Chr26. The fate of LSU_18 in *M. aruncella* is a remarkable case of chromosome instability. While it was intact in the trichopteran species as ell as in all heteroneuran species it contributed to 28 of the 31 chromosomes of *M. aruncella*, almost always in small fractions of the unit, 1 - 9 of 129 markers. The only major contribution, 44 of the 129 markers, was made to the Z chromosome. where it joined the original Z chromosome block, LSU_31.

#### Discussion

### Lepidopteran Synteny Units and chromosome number

Here we show that lepidopteran genomes are composed of well-conserved blocks of synteny which we term Lepidopteran Synteny Units or LSUs. There are 31 of them in the lepidopteran clade Heteroneura which contains more than 99% of the ∼160,000 butterfly and moth species. Each LSU defines the composition of a single chromosome in the representative species with karyotypes of n = 31 chromosomes. In those with less chromosomes, some chromosomes are composed of two or three LSUs combined together (see Figures 2 and 3). The analysis of the trichopteran *L. marmoratus* showed that 24 of the LSUs already existed as chromosomal entities before Lepidoptera and Trichoptera lineages split, some 230 Mya (Wiegmann et al. 2009). This estimate of age may indeed be too conservative. According to Kawahara et al. (2019) the oldest members of crown Lepidoptera lived ∼300 Mya. The full set of LSUs came into being after Micropterigidae branched off from the main stem of Lepidoptera and before the diversification of Heteroneura, about 172 Mya (Wahlberg et al. 2013), or ∼230 Mya according to Kawahara et al. (2019).

The modal number of chromosomes in Lepidoptera is 30 - 31 (Robinson 1971). The occurrence of species with n = 31 chromosomes, each represented by a single LSU, in Adelidae, Yponomeutidae, Cossidae, Plutellidae, Nymphalidae, Geometridae, and Noctuidae confirms that n = 31 is the basal karyotype in the Heteroneura clade of Lepidoptera.

Karyotypes with a different chromosome number are derived from it. The ancestral chromosome number at the root of Lepidoptera, however, cannot yet be deduced. Although the micropterigid *M. aruncella* has n = 31 chromosomes, the same number, the chromosomes are not the same. In this species as well as in the trichopteran *L. marmoratus*, LSU_2 is found split into two chromosomes but different LSUs are fused (see Table S3).

### Sex chromosomes

The Z chromosomes of all species studied here share the same LSU, LSU_31. This means that Lepidoptera and Trichoptera have the same Z chromosome, as noticed earlier by Fraїsse et al. (2017). In *B. adustella*, the ancestral Z chromosome, represented by LSU_31, is fused with an autosome, represented by LSU_21 and in *M. aruncella* the original Z chromosome is enlarged by an originally autosomal chunk from LSU_18. In a few of the selected species, the female restricted W chromosomes were sequenced, however they are not included here as no BUSCO hits were recorded on them.

### Rarrangements within and between LSUs

We have shown that LSUs are highly conserved across the Heteroneura and large rearrangements between LSUs are not found within the chosen species spanning this group. Gains and losses of LSUs therefore reflect a more subtle pattern of ‘leakage’ whereby individual LSUs are present in some species and absent in others. However this pattern is not ubiquitous. A few ‘hot spots’ of interchromosomal exchange were observed within the sampled genomes. LSU_18 in *M. aruncella* is nearly pulverized except for 44 of the 129 markers enlarging the Z chromosome of this species, 85 of the 129 markers were contributed to various other chromosomes. Not quite as dramatic, LSU_28 in *P. xylostella* contributed 41 of 55 markers to Chr_31 and 14 to other chromosomes. It would be interesting to learn what causes the instability in these chromosomes.

### Synteny in other taxa and future applications

Blocks of conserved synteny have been reported from studies in fungi (e.g. Li et al. 2022), plants (e.g. Phan et al. 2007) and animals (e.g. Hui et al. 2012; Simakov et al. 2022).

However, this report is the first systematic attempt to document synteny genome-wide in butterflies and moths. It also puts forward a simple application of BUSCO analysis as a robust method of defining macrosynteny in any taxa. The conserved synteny blocks, described by the LSU markers, are surprisingly well conserved in the Heteroneura, representing ∼200 My of conservation. In contrast, macrosynteny decayed very fast, within ∼100 My, in budding yeast species of the subphylum Saccharomycotina (Li et al. 2022). And it decayed similarly fast in nematodes: Coghlan and Wolfe (2002) detected 2.3 - 5.4 reciprocal translocations / My in comparisons of the *Caenorhabditis elegans* and *C. briggsae* genomes.

In terms of future applications, LSUs are easily derived from BUSCO analysis of lepidopteran target genomes and are a simple means to describe the architecture of lepidopteran genomes and their chromosomal evolution. The more complex karyotypes of some lycaenid (blue butterflies) (Lukhtanov 2015) or pierid (white) butterflies (Šíchová et al. 2016, Höög et al. 2023) are obvious targets for further analysis. BUSCO analysis with LSUs also offers another practical use: to quickly determine places of interest like breakpoints in chromosomal rearrangements for a subsequent detailed analysis of the nature of such breakpoints or fusion points at the nucleotide level. Finally, we expect that this type of homology-based analysis can also be applied to other groups of animals or plants where similar chromosome-wide blocks of syntenic markers can readily be established. Understanding and accurately describing macrosynteny is key if we are ever to understand why chromosome number varies so considerably in both animals and plants.

## Abbreviations

Mya: million years ago
My: million years
LSU: Lepidopteran Synteny Unit
BUSCO: Benchmarking Universal Single-Copy Orthologs

## Acknowledgements

A part of this research was supported by JSPS to KS (19K06067).

## Figure captions

## Supplementary material

**Table S1** Genomes used in this study.

Species, family, accession number, chromosome number

**Table S2** LSU-specific lepidoptera_odb10 subsets, ordered according to chromosome position in *Melitaea cinxia*

**Table S3** Conservation of LSUs. The table lists the number of markers from a specific LSU found on a specific chromosome compared with that on the whole genome (chromosome / genome).

## Availability of data and materials

All genome data are available in public data banks with the acc. nos. given in the manuscript. Any additional details such as Java code will gladly be provided upon request.

